# Inhibitors of L-type calcium channels show therapeutic potential for treating SARS-CoV-2 infections by preventing virus entry and spread

**DOI:** 10.1101/2020.07.21.214577

**Authors:** Marco R. Straus, Miya K. Bidon, Tiffany Tang, Javier A. Jaimes, Gary R. Whittaker, Susan Daniel

**Author notes:** Corresponding authors: Gary Whittaker, Susan Daniel.

## Abstract

COVID-19 is caused by a novel coronavirus, severe acute respiratory syndrome coronavirus (CoV)-2 (SARS-CoV-2). The virus is responsible for an ongoing pandemic and concomitant public health crisis around the world. While vaccine development is proving to be highly successful, parallel drug development approaches are also critical in the response to SARS-CoV-2 and other emerging viruses. Coronaviruses require Ca^2+^ ions for host cell entry and we have previously shown that Ca^2+^ modulates the interaction of the viral fusion peptide with host cell membranes. In an attempt to accelerate drug development, we tested a panel of L-type calcium channel blocker (CCB) drugs currently developed for other conditions, to determine whether they would inhibit SARS-CoV-2 infection in cell culture. All the CCBs tested showed varying degrees of inhibition, with felodipine and nifedipine strongly limiting SARS-CoV-2 entry and infection in epithelial lung cells at concentrations where cell toxicity was minimal. Further studies with pseudo-typed particles displaying the SARS-CoV-2 spike protein suggested that inhibition occurs at the level of virus entry. Overall, our data suggest that certain CCBs have potential to treat SARS-CoV-2 infections and are worthy of further examination for possible treatment of COVID-19.

## Introduction

Coronaviruses are major zoonotic pathogens that cause respiratory and/or enteric tract infections in a variety of species, including humans. Most of the coronaviruses that are pathogenic to humans cause only mild common cold-like disease symptoms^1,2^. However, currently a COVID-19 pandemic, caused by the severe acute respiratory syndrome (SARS)-CoV-2, poses a dramatic risk to public health worldwide^3^. The virus was first identified in December 2019 in Wuhan, China and spread rapidly across the globe^4^. To date, several vaccines have been approved for emergency use^5^, however, the worldwide demand for these vaccines faces major difficulties due to the current limitations in the vaccine production and distribution, which reiterates the need for alternative therapeutic strategies to mitigate the clinical impact of the disease. Current antiviral treatment for SARS-CoV-2 is limited to a viral polymerase inhibitor, remdesivir, used under emergency use authorization; however, this has proved to be of only limited benefit in COVID-19 patients^6^. Thus, there remains an important need to identify and to provide additional drugs that can treat coronavirus infections.

The entry of the virus into the host cells is a promising target for a potential therapeutic, as it is a crucial step in the viral life cycle^7^. Possible intervention steps include ACE-2 blockers^8^, host cell protease inhibitors such as camostat^9^ and inhibitors of virus fusion^10,11^, as well as monoclonal antibodies directed toward spike^12^. Previous studies have revealed that coronaviruses, such as SARS-CoV and MERS-CoV utilize calcium ions (Ca^2+^) for viral entry, via coordination of the ion by amino acid residues within the conserved fusion peptide of the viral spike (S) protein^13,14^. In these cases, depleting either intracellular and/or extracellular Ca^2+^ with chelating agents results in full or partial inhibition of viral entry and fusion^13,14^. This suggests a novel way to inhibit coronavirus fusion and entry more broadly. Recently, the impact of calcium on SARS-CoV-2 fusion was examined through biophysical studies^15,16^, which showed broad functional similarity between SARS-CoV and SARS-CoV-2 with regard to Ca^2+^ interactions.

The important role of Ca^2+^ in the entry of these coronaviruses prompted us to explore whether Ca^2+^ channel blocker (CCB) drugs inhibit SARS-CoV-2 in cell culture and have the potential to be used therapeutically. In this study, we chose five drugs from different classes that all inhibit high voltage-activated Ca^2+^ channels of the L-type. We selected the dihydropyridines amlodipine, nifedipine and felodipine; the phenylalkylamine verapamil and the benzothiazepine diltiazem. These five drugs are primarily used to treat cardiovascular diseases, including hypertension. In addition, we also tested a Ca^2+^ chelator drug: diethylenetriaminepentaacetic acid (DTPA), which is used to treat radioactive contamination of internal organs. We wanted to also include Ca^2+^ chelators, as previous studies have shown that EGTA and BAPTA-AM (intracellular chelator) successfully inhibited entry in other coronaviruses^13,14^. Our studies reveal the potential for developing new treatment modalities for COVID-19 through the use of L-type calcium channels blockers.

## Results and Discussion

While many aspects of SARS-CoV-2 entry remain unresolved, Vero E6 cells are widely used as cell line for propagation of the virus. As an initial screen of our drug panel, we infected Vero E6 cells at a relatively low MOI (0.1 infectious units per cell) in order to monitor virus propagation. We carried out a dose-response study, by adding four different concentrations of each compound one hour post infection (10 µM, 50 µM, 100 µM and 500 µM). Changes in viral propagation were then assessed after 24 h by TCID_50_ assay. We also monitored cell viability under equivalent conditions. This data is shown in Figure 1. For amlodipine, we observed a 5-log reduction of viral growth at 50 µM and we were not able to detect any viral titer at a concentration of 100 µM (Figure 1A). However, amlodipine also exhibited a 25% reduction of cell viability at a concentration of 50 µM, which increased to 90% at 100 µM and 500 µM (Figure 1A). In comparison, nifedipine reduced viral titers by 5 logs at 500 µM and showed about 15% cytotoxicity (Figure 1B). Felodipine completely suppressed growth of SARS-CoV-2 at a concentration of 50 µM with no statistically significant cytotoxicity (Figure 1C). 100 µM verapamil reduced the viral titers by 50% compared to the untreated control without a cytotoxic effect (Figure 1D). Diltiazem reduced viral growth at 500 µM but also compromised cell viability at this concentration and DTPA showed no effect Figure 1E and 1F).

**Figure 1:**
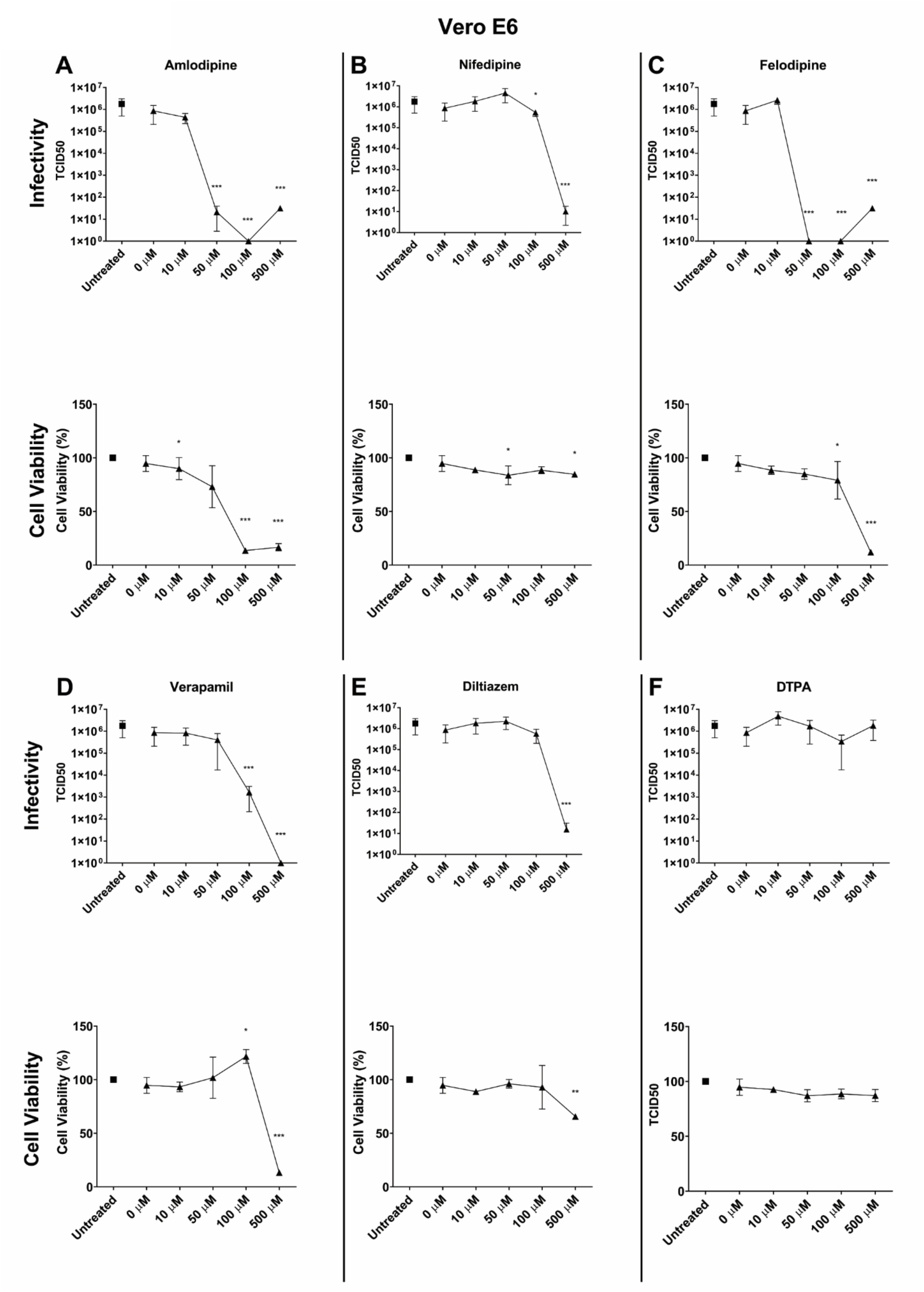
Inhibitory effect of five CCBs and the Ca^2+^ chelator DTPA on SARS-CoV-2 infection and correlation to cell viability in Vero E6 cells. **(Infectivity A-E)** Vero E6 epithelial kidney cells were infected with SARS-CoV-2 isolate USA-WA1/2020 at a MOI of 0.1 for 24 hours. The CCBs amlodipine, nifedipine, felodipine, verapamil and diltiazem and the chelator DTPA were added to the cells at the indicated concentrations immediately after the virus. 0 µM sample contained only DMSO which was used as solvent for the drugs. TCID_50_ assays were performed with growth supernatants and calculated according to the Reed-Muench method^40^. No bar means no virus was detected at the respective concentration. Error bars represent standard deviations (n = 3). Asterisks indicate statistical significance compared to the untreated control. Statistical analysis was performed using an unpaired Student’s t-test. ^*^ = P > 0.05, ^**^ = P > 0.01, ^***^ = P > 0.001. **(Cell viability A-F)** Vero E6 epithelial kidney cells were treated with the indicated concentrations of amlodipine, nifedipine, felodipine, verapamil, diltiazem and DTPA for 24 hours. After 24 hours cell viability was measured using 3-(4,5-dimethylthiazol-2-yl)-2,5-diphenyltetrazolium bromide (MTT). Cell viability was determined by normalizing absorbance from the sample well by the average absorbance of untreated wells. As for the infections, 0 µM sample contained only DMSO which was used as solvent for the drugs. Error bars represent standard deviations (n = 3). Asterisks indicate statistical significance compared to the untreated control. Statistical analysis was performed using an unpaired Student’s t-test. ^*^ = P < 0.05, ^**^ = P < 0.01, ^***^ = P < 0.001.

While Vero E6 cells are a good model system for SARS-CoV-2 infections, the cell line is derived from epithelial kidney cells. As such, these cells may not represent the virus-drug-cell interactions in the respiratory tract and may show differences in entry pathway. Therefore, we repeated our experiments in the human lung epithelial cell line Calu-3 (Figure 2). The results for amlodipine were comparable to that found in Vero E6 cells (Figure 1A and 2A). Nifedipine and felodipine, however, significantly inhibited viral growth at lower concentrations compared to Vero E6 cells (Figure 2B and 2C). Nifedipine reduced the viral titers by 1.5 logs at a concentration of 100 µM and no virus was detectable at 500 µM while cytotoxicity was moderate (Figure 2B). Felodipine diminished SARS-CoV-2 growth by approximately half at 10 µM and at 50 µM no virus was detected with no cytotoxic effect on the cells (Figure 2C). In contrast, verapamil had a weakened inhibitory effect at 100 µM compared to Vero E6 cells (Figure 2D). As in Vero E6 cells, 100 µM verapamil fully suppressed viral growth but also compromised cell viability by about 90%. We found very modest infection inhibition for diltiazem and DTPA at non-cytotoxic concentrations of 500 µM and 100 µM, respectively (Figure 2E and 2F).

**Figure 2:**
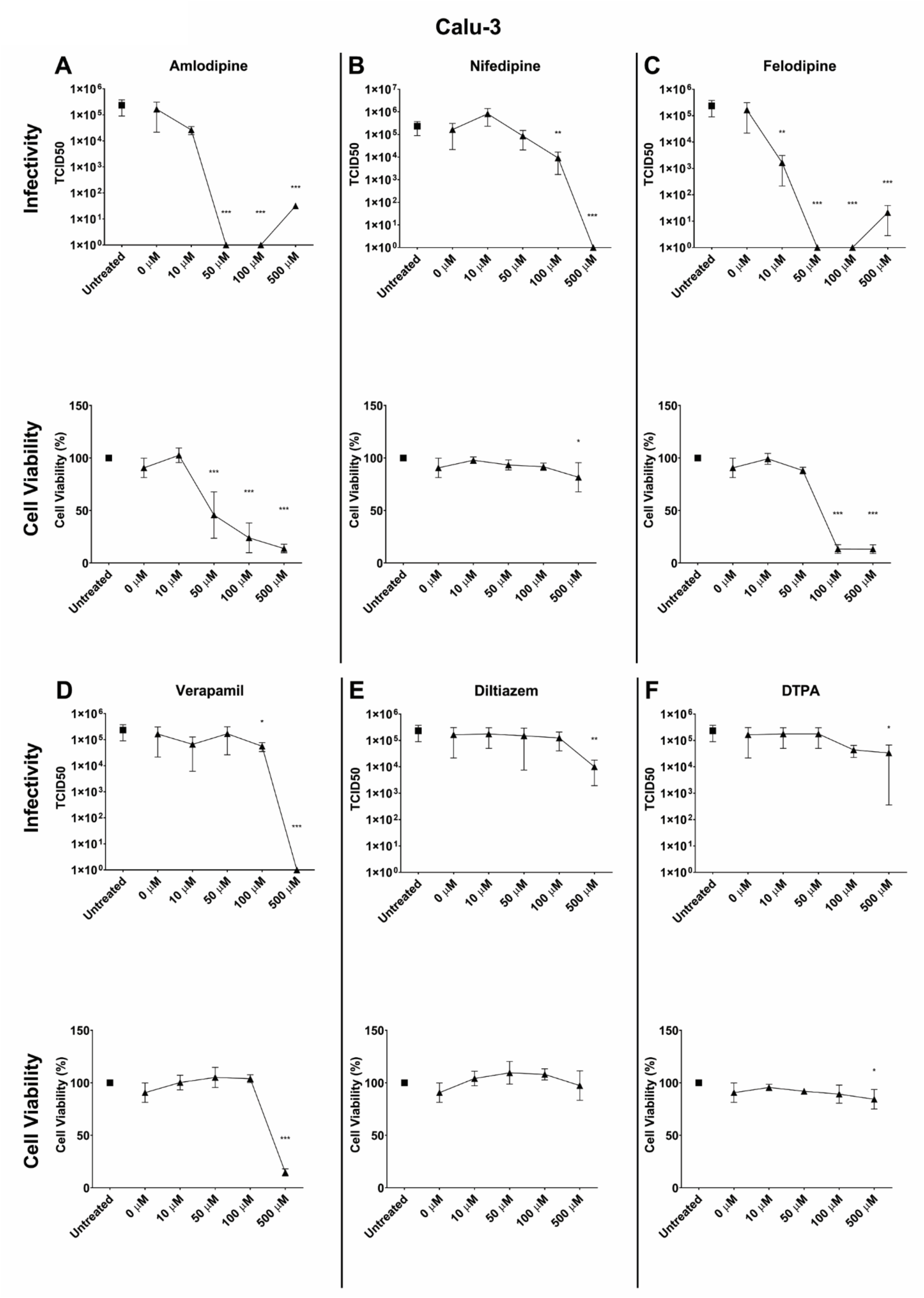
Inhibitory effect of five CCBs and the Ca^2+^ chelator DTPA on SARS-CoV-2 infection and correlation to cell viability in Calu-3 cells. Conditions for **(Infectivity A-F)** and **(Cell viability A-F)** in Calu-3 epithelial lung cells as described in figure legend Figure 1. Error bars represent standard deviations (n = 3). Asterisks indicate statistical significance compared to the untreated control. Statistical analysis was performed using an unpaired Student’s t-test. ^*^ = P < 0.05, ^**^= P < 0.01, ^***^ = P < 0.001.

A possible explanation for the observed difference could be found in varying expression levels of L-type calcium channels in Vero E6 and Calu-3 cells and we will pursue this question in our follow-up study. In this context it is noteworthy that L-type CCBs are categorized in different chemical classes. Felodipine and nifedipine as well as amlodipine belong to the class of dihydropyridines (DHPs) while diltiazem is a benzothiazepine (BZPs) and verapamil is a phenylalkamines (PAAs). All three classes bind to the α_1_ subunit of the L-type calcium channel but their binding affinity is state dependent^17–19^. DHPs preferentially bind to the channel in its inactive state when there is an increased membrane potential^18^. In contrast, PAAs and BZPs have a higher affinity to stimulated L-type calcium channels at a lower membrane potential^19^. Hence, it is conceivable that SARS-CoV-2 infection impacts the membrane potential within cells and therefore influences the affinity of the different CCBs to the α_1_ subunit of L-type calcium channels.

Furthermore, both diltiazem and verapamil were shown to inhibit calcium signaling from nicotinic acid adenine dinucleotide phosphate (NAADP) triggering^20^, which are major activators of two pore calcium channels (TPCC)^21^. It was shown that using either diltiazem or verapamil to inhibit TPCC function was effective in inhibiting Ebola virus infection^22^. However, our results demonstrate modest inhibition from either diltiazem or verapamil when compared with the inhibition from felodipine, amlodipine, or nifedipine treatment. This suggests that calcium signaling resulting in proper TPCC function may not be critical for SARS-CoV-2 infection.

With amlodipine, felodipine and nifedipine being the most promising candidates for suppression of SARS-CoV-2 growth in vivo, we determined the selectivity index (SI) (also referred to as TI, therapeutic index) of these compounds. The SI provides a measure how safe a given drug would be for the treatment of SARS-CoV-2 infections *in vivo*. To calculate the SI, we determined the concentration at which they inhibit 50% of the viral growth (EC_50_) and the concentration at which each drug exerts 50% cytotoxicity (CC_50_). Dividing CC_50_/EC_50_ results in the SI, and the higher the SI the more selective the drug is against the pathogen. However, it must be emphasized that while the SI determined here serves as an indicator and provides important information to consider for human application, the final therapeutic efficacy in humans may differ and potential side effects of the drugs cannot be taken into account. We used Calu-3 cells as the more relevant cell line compared to Vero cells. Cells were infected with SARS-CoV-2 and different concentrations of the respective drugs were applied (Figure S1). The obtained EC_50_ values were 10.36 µM for amlodipine, 0.01255 µM for felodipine and 20.47 µM for nifedipine (Figure S1A - C, Table 1). We determined the CC_50_s in a similar fashion using a cytotoxicity assay as described above resulting in CC_50_ values of 27.85 µM for amlodipine and 122.8 µM for felodipine (Figure S1A and B). For nifedipine the highest tested concentration of 2 mM did not significantly impair cell viability (Figure S1C, Table 1). The obtained SI values were 2.69 for amlodipine and 9,784.86 for felodipine (Table 1). Because nifedipine did not show a significant cytotoxic effect at approx. 7x the concentration of the most efficacious antiviral concentration of 300 µM a SI value could not be reached. Hence, all three drugs seem to be highly selective against SARS-CoV-2 with felodipine and nifedipine being the most promising candidates because of their high SI score and very low cytotoxicity, respectively.

**Table 1:**
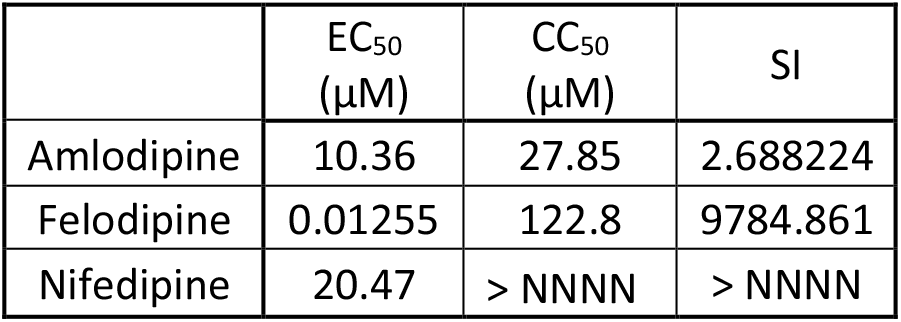
EC_50_, CC_50_ and Selectivity Index (SI) of Amlodipine, Felodipine and Nifedipine. EC_50_ and CC_50_ were determined in Calu-3 cells. SI = CC_50_/EC_50_.

| | $\text{EC}_{50}$<br>($\mu\text{M}$) | $\text{CC}_{50}$<br>($\mu\text{M}$) | SI |
| --- | --- | --- | --- |
| Amlodipine | 10.36 | 27.85 | 2.688224 |
| Felodipine | 0.01255 | 122.8 | 9784.861 |
| Nifedipine | 20.47 | > NNNN | > NNNN |

It is important to emphasize that felodipine and nifedipine work with higher efficiency in Calu-3 cells compared to Vero E6 cells (Figure 1 and 2). In this context, it is noteworthy that CCBs are believed to affect airway smooth muscle cells, which rely on L-type calcium channels for their contraction^23–25^. Recently, the same CCBs tested here were shown to have a beneficial effect on the lung function of asthma patients, providing evidence that CCBs act in the lungs^26^. More evidence that interference with the cellular Ca^2+^ balance suppresses viral growth comes from nebulizers used for asthma medications. Nebulizers are supplemented with EDTA as a preservative, which may help to deplete extracellular Ca^2+ 27^. and may provide a clue as to why asthma is not on the top ten list of chronic health problems of people who died of COVID-19.

While these results are promising, it is unclear how the efficacious doses of amlodipine, nifedipine and felodipine found here would translate into clinical use in human patients to treat COVID-19 infections. Nevertheless, a recent report suggests that amlodipine and nifedipine reduce mortality and the risk for intubation in COVID-19 patients^28^ and hence, provides an encouraging example that CCBs could be a viable option to fight COVID-19 infections.

The inhibition differences observed here in these two cell lines, combined with previous work from our labs, suggest that inhibition may occur at the level of host cell entry. To further investigate whether CCB-mediated inhibition of SARS-CoV-2 infectivity affects viral host cell entry, we utilized murine leukemia virus (MLV)-based pseudo particles (PPs), to infect both VeroE6 and Calu-3 cells. The pseudoparticles are decorated with the SARS-CoV-2 S protein^29^ and carry a genome encoding for luciferase, so that infected cells produce luciferase upon transduction, which is quantifiable. These virion surrogates allow for only one infection cycle without intracellular replication and thus, inhibition of PP infection would suggest that the CCBs affect host cell entry. As a control, PPs harboring the SARS-CoV-2 S protein were shown to be several orders of magnitude more infectious than their no envelope (Δenv) counterpart (Figure S2).

To target virus entry specifically, we pretreated all cells with the drugs for one hour before adding PPs carrying the SARS-CoV-2 S protein (Figure 3). We repeated the cytotoxicity studies in which cells were incubated with the drugs for 72 hours to match the PP experiments (Figure S3) and the trends were similar to the results in Figure 2, except that Diltiazem at 500 µM and DTPA after 50 µM exhibited greater cytotoxicity after 72-hour incubation in both the Vero E6 and Calu-3 cell line. Amlodipine suppressed PP infection at 50 µM in both, Vero E6 and Calu-3 cells (Figure 3). Nifedipine inhibited PP infection in Vero E6 and Calu-3 cells with 500 µM having the strongest effect. Felodipine inhibited PP entry in both cell lines at 50 µM. Consistent with what we found in the live virus infection experiments presented above, verapamil suppressed PP entry at 50 µM in Calu-3 and at 100 µM in Vero E6 cells. In contrast to our observations with SARS-CoV-2 live virus infections, diltiazem and DTPA also inhibited PP infections (Figure 1, 2 and 3). Diltiazem inhibited PP infection in Calu-3 starting at 50 µM, and in Vero E6, at 500 µM, however we note that diltiazem is cytotoxic to both cell lines at 500 µM. Although we observe that DTPA inhibits PP entry, we also note that DTPA is cytotoxic to the cells after 72 hours at 50 µM (Vero E6) and 10 µM (Calu-30), and so we believe that the inhibition could be attributed to the cytotoxicity of DTPA.

**Figure 3:**
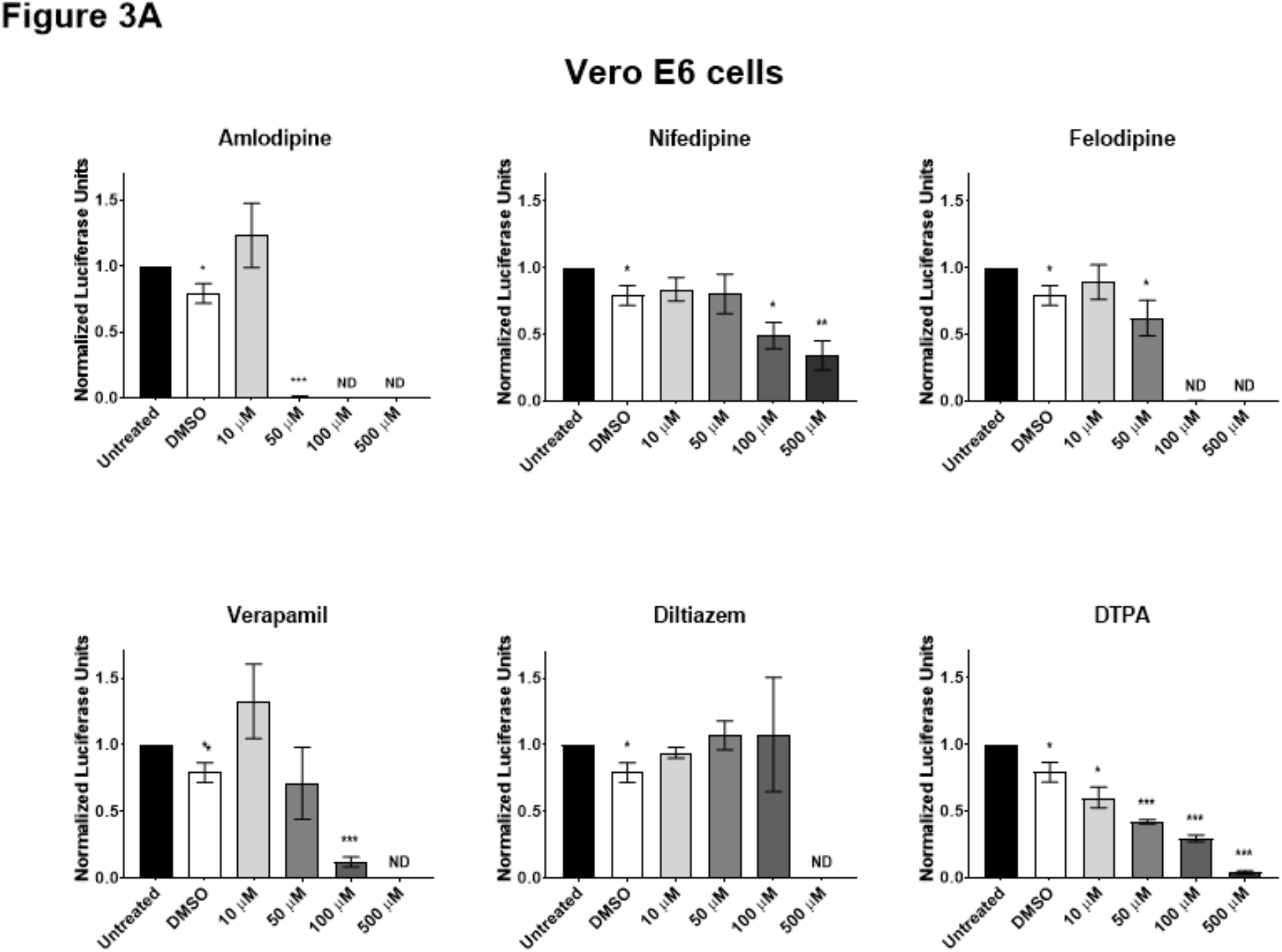
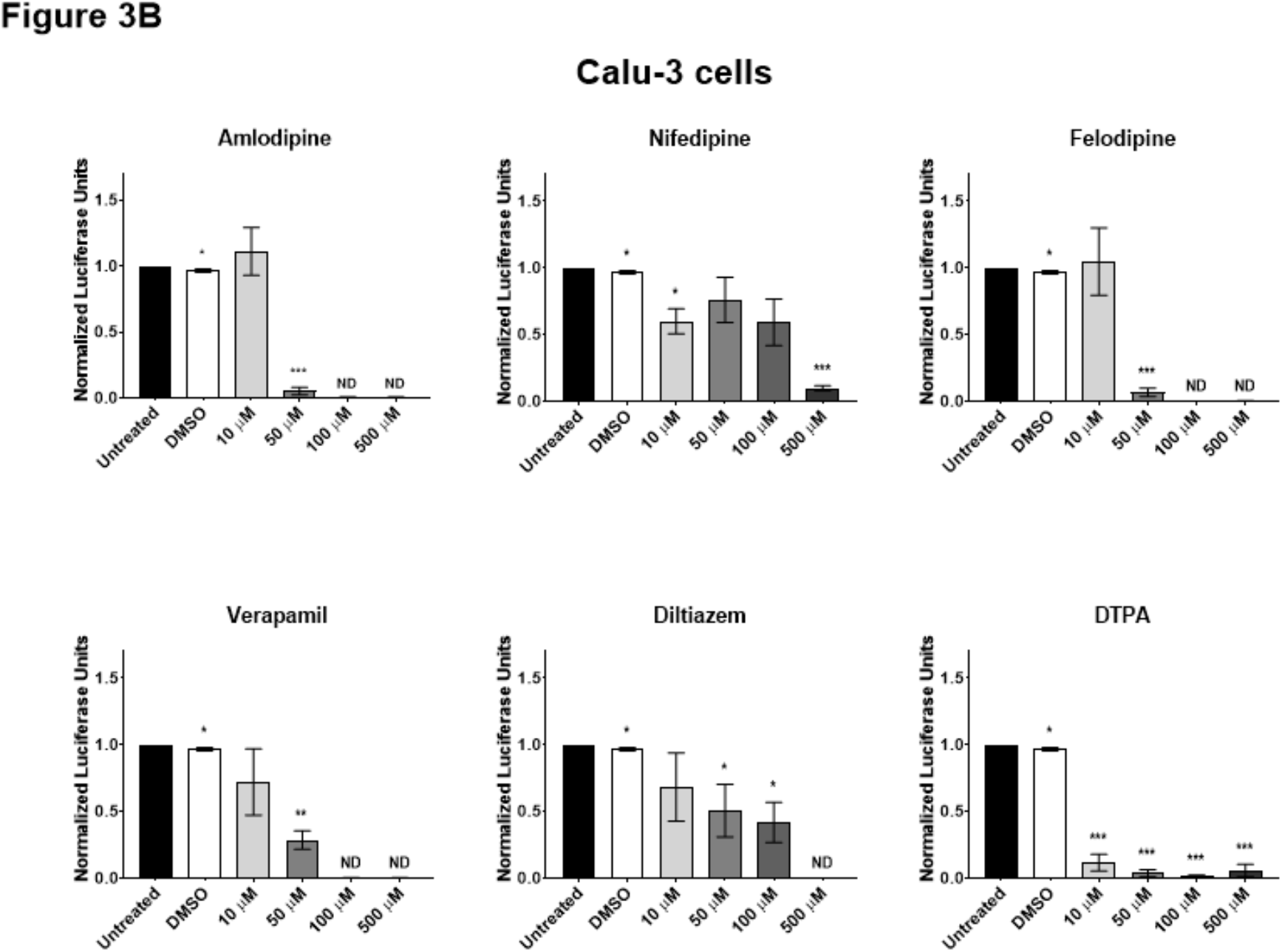
Pseudo particle assays of SARS-CoV-2 S in Vero E6 and Calu-3 cells. (A) Vero E6 epithelial kidney cells and (B) Calu-3 epithelial lung cells were pre-treated with CCBs and DTPA for one hour prior to infection with PPs carrying SARS-CoV-2 S. After 24 hours growth medium was changed and replenished with the drugs. 72 hours after infection the luciferase activity was assessed. Infectivity was normalized to the untreated sample. Error bars represent standard deviations (n = 3). Asterisks indicate statistical significance compared to the untreated control. Statistical analysis was performed using an unpaired Student’s t-test. ^*^ = P < 0.05, ^**^ = P < 0.01, ^***^ = P < 0.001.

In Vero cells, CoVs are believed to preferentially enter the host cells via the endocytic pathway^7,30^. In contrast, in Calu-3 cells, SARS-CoV-2 spike is predominantly activated at the plasma membrane ^7,31^; however specific entry pathways and fusion location remain unresolved. There is evidence from previous reports that CCBs inhibit other viral infections at the cell entry level, even for viruses without an apparent Ca^2+^ ion-binding to the fusion peptide. Influenza virus infection, for example, triggers an influx of Ca^2+^ that assists the endocytic uptake of the virus, with treatment by verapamil or diltiazem inhibiting viral infection^32–34^. New World hemorrhagic fever arenaviruses such as Junin virus are reportedly sensitive to CCBs and treatment with CCBs blocks the entry of the virus into the host cell^35^. CCBs were also shown to interfere with viral replication of several other viruses (*e*.*g*., Japanese Encephalitis virus, Zika virus, Dengue virus)^36^. A recently published study observed SARS-CoV-2 inhibitory activity when cells were treated with amlodipine, and attributed it to the fact that amlodipine upregulates cholesterol levels which may block SARS-CoV-2 infection^37^. Thus, the exact mechanism of how CCBs suppress these viruses, as well as SARS-CoV-2 infection, needs to be more fully addressed. While our results point to an interference of SARS-CoV-2 at host cell entry, it is conceivable that CCB-mediated inhibition of viral spread occurs at other stages as well, including viral release. It will require further studies to determine whether CCBs interfere directly with virus-cell fusion, how these drugs may affect the different fusion pathways and to explore whether CCBs may inhibit other cell functions that subsequently lead to inhibition of host cell entry.

Regardless of the exact mechanism, these examples demonstrate that CCBs exhibit an antiviral efficacy against a broad range of viral pathogens relevant for public health. They also show the broad requirement of Ca^2+^ ions for viral propagation, supported by other studies that report a Ca^2+^ requirement for major human pathogens like Ebola virus and Rubella virus^38,39^. Therefore, CCBs may represent a novel class of antiviral therapeutics against a broad range of major viral diseases that warrant further clinical studies, but especially to address the current crisis of COVID-19.

In summary, the results described above are promising and particularly felodipine and nifedipine may represent a viable treatment option against COVID-19. However, we emphasize that it is unclear how the efficacious doses found here and the SI score would translate into clinical use in human patients to treat COVID-19 infections. As a next step, a meta-analysis of patient data taking CCBs medication and the relation to the outcome and severity of COVID-19 (e.g., hospitalization, intubation etc.) in these patients may provide further insights about the efficacy of CCBs to inhibit SARS-CoV-2 infections in humans. This is the focus of our future studies.

## Methods

### Cells and reagents

Vero E6 and Calu-3 cells were obtained from the American Type Culture Collection (ATCC). Cells were maintained in Dulbecco’s modified Eagle medium (DMEM) (Cellgro) supplemented with 25 mM HEPES (Cellgro) and 10% HyClone FetalClone II (GE) at 37° C and 5% CO_2_. For virus infections, cells were grown in Eagle’s Minimum Essential Medium (EMEM) (Cellgro) supplemented with 4% heat inactivated fetal bovine serum (FBS) (Gibco) at 37° C and 5% CO_2_.

The SARS-CoV-2 isolate USA-WA1/2020 was obtained from the Biological and Emerging Infections Resources Program (BEI Resources). Amlodipine, nifedipine, felodipine, verapamil, diltiazem and DTPA were purchased from Sigma Aldrich. 3-(4,5-dimethylthiazol-2-yl)-2,5-diphenyltetrazolium bromide (MMT) was obtained from Thermo Fisher. Crystal violet was purchased from VWR. pCDNA3.1/SARS-CoV-2 S Wuhan Hu-1 was generously provided by Dr. David Veesler.

### SARS-CoV-2 infections and TCID_50_ assays

Vero E6 and Calu-3 cells were grown to confluency under biosafety level-2 (BSL-2) conditions. Cells were then transferred to the BSL-3 lab, washed with DBPS and a volume of 200 µL infection media with SARS-CoV-2 at a MOI of 0.1 was added to the cells. Cells were then incubated at 37° C and 5% CO_2_ for 1 hour on a rocker. Cells were then supplemented with 800 µL infection media (EMEM + 4% Fetal Bovine Serum) and appropriate concentrations of each drug were added. Amlodipine, nifedipine, felodipine and verapamil were dissolved in DMSO. Therefore, DMSO was used as a control at the same volume that was applied for the highest drug concentration. After 24 hours, the supernatants were harvested and stored at -80° C.

For the TCID_50_, Vero E6 cells were grown to confluency in 96 well plates and serial dilution of the supernatants were prepared. Undiluted and diluted samples were then added to the Vero E6 cells and grown for 72 hours at 37° C and 5% CO_2_. After 72 hours supernatants were aspirated, and cells were fixed with 4% paraformaldehyde. Cells were then stained with 0.5% crystal violet and subsequently washed with dH_2_O. Wells were scored and analyzed for living or dead cells according to the Reed-Muench method^40^.

### Cytotoxicity assay

The cytotoxicity of the calcium ion blocking drugs (amlodipine, nifedipine, felodipine, diltiazem) and calcium chelator (DTPA) on VeroE6 and Calu-3 cells were determined by an 3-(4,5-dimethylthiazol-2-yl)-2,5-diphenyltetrazolium bromide (MTT) assay. A total of 5 × 10^5^ VeroE6 cells/well, and 6.7 × 10^5^ Calu-3 cells/well were incubated in the presence of the six calcium blocking drugs and calcium chelator at the concentrations of 10, 50, 100, and 500 µM for 24 or 72 hour at 37°C C and 5% CO_2_ in a CO_2_ incubator. After 24 or 72 hours incubation, cells were treated with MTT solution (5 mg/mL) and incubated for 4 hours at 37°C and 5% CO_2_ with rocking to allow purple formazan crystals to form. 50 µL DMSO was added into each well to dissolve the crystals. The absorbance was measured at 540 nm in a microplate reader (Bio-Tek Instrument Co., WA, USA).

### Pseudo particle production and infections

Murine leukemia virus (MLV) pseudo particles production and infections were performed as previously described with minor modification^14^. For PP production, HEK 293T cells were transfected with pTG-luc luciferase reporter, pCMV-MLV-gagpol, and pCDNA3.1/SARS-CoV-2 S Wuhan Hu-1 to generate SARS-CoV-2 S carrying PPs. Supernatants were harvested 72 hours post transfection and stored at -80°C. For infections, cells were seeded in 96-well plates and DMEM containing the different CCBs and DTPA was added and incubated for 1 hour. After one hour 50 µL PPs were added. After 24 hours, cells were replenished with DMEM containing the individual drugs at the appropriate concentration and incubated for another 48 hours to allow expression of luciferin at sufficient levels for detection. After lysis, luciferase activity was measured using a GloMax Navigator Microplate Luminometer (Promega).

### Statistical analysis

Individual comparisons between the treatments and the control (DMSO treated) were performed for both live virus and pseudo particle assays. The results were analyzed using an unpaired Student’s t-test using Microsoft Excel and GraphPad Prism 8 to make individual comparisons.

## Acknowledgments

This projected was funded by Fast Grant, Mercatus Center. We would like to thank Paul Jeannette for his support for all the BSL-3 work and Dr. Luis Schang, Dr. Nihal Altan-Bonnet, Dr. Fernando Martinez, Dr. Bruce Kornreich and Dr. Hanno Andreas Ludewig for their critical input. TT is supported by the National Science Foundation Graduate Research Fellowship Program under Grant No. DGE-1650441.

## Supporting Information

Supporting information available: Determination of EC_50_ and CC_50_ of Amlodipine, Felodipine, and Nifedipine in Calu-3 cells (**Figure S1**); Characterization of the MLVpp system (**Figure S2**); 72-hr cell viability study (**Figure S3**).

---

**Figure S1:**
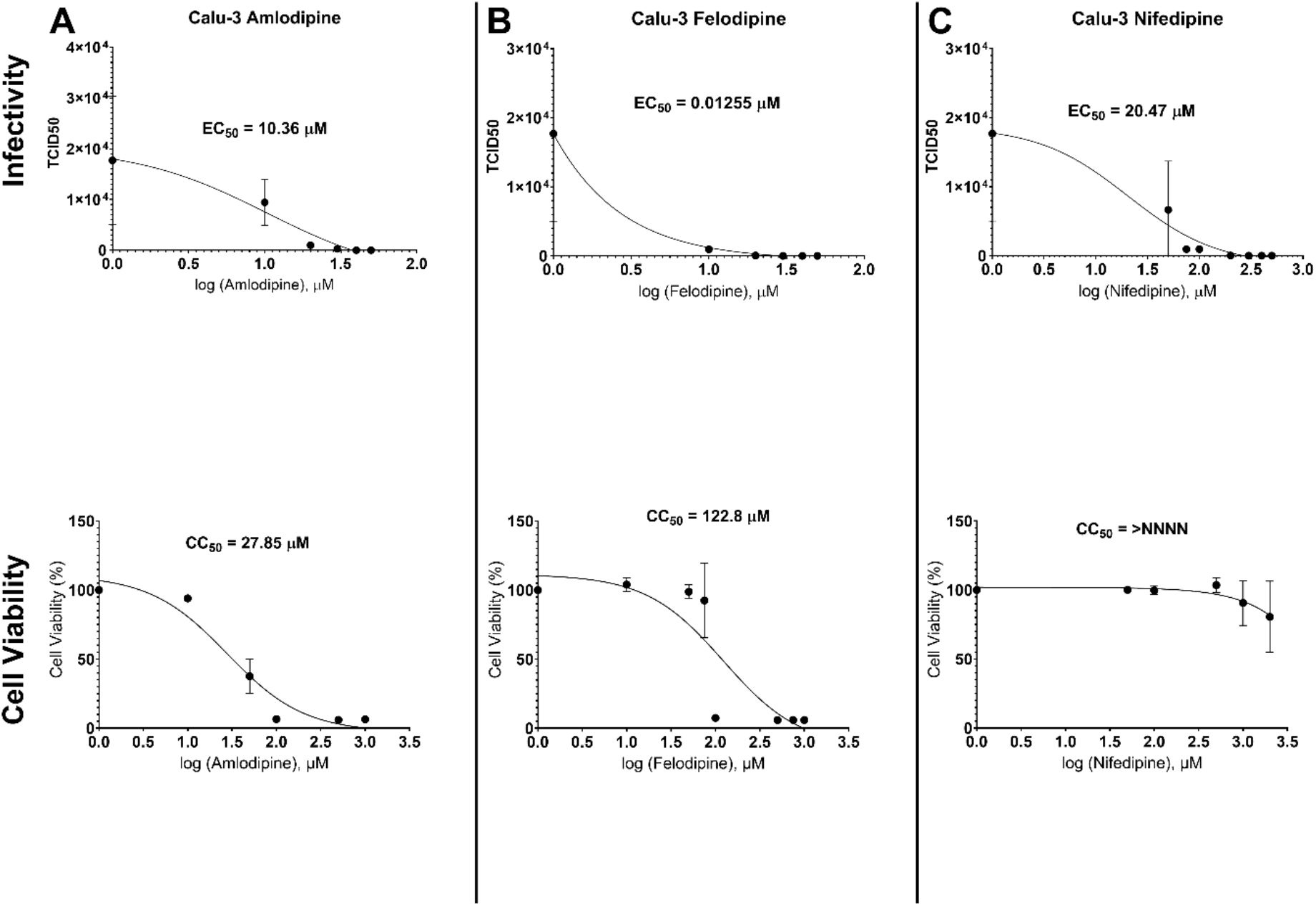
Determination of EC_50_ and CC_50_ of Amlodipine, Felodipine and Nifedipine in Calu-3 cells. Calu-3 epithelial lung cells cells were infected with SARS-CoV-2 isolate USA-WA1/2020 at a MOI of 0.1 for 24 hours. The CCBs amlodipine (A) and felodipine (B) were added to the cells at the 0, 10, 20, 30, 40 and 50 µM immediately after the virus. Nifedipine (C) was added at concentrations of 50, 75, 100, 200, 300, 400 and 500 µM. TCID_50_ assays were performed with growth supernatants and calculated according to the Reed-Muench method^40^. Cell viability was measured using 3-(4,5-dimethylthiazol-2-yl)-2,5-diphenyltetrazolium bromide (MTT). Cell viability was determined by normalizing absorbance from the sample well by the average absorbance of untreated wells. Drug concentrations were converted into log10 numbers and plotted on the x-axis. EC_50_ and CC_50_ values were calculated using Graphpad Prism 8 software. Error bars represent standard deviations (n = 3).

**Figure S2:**
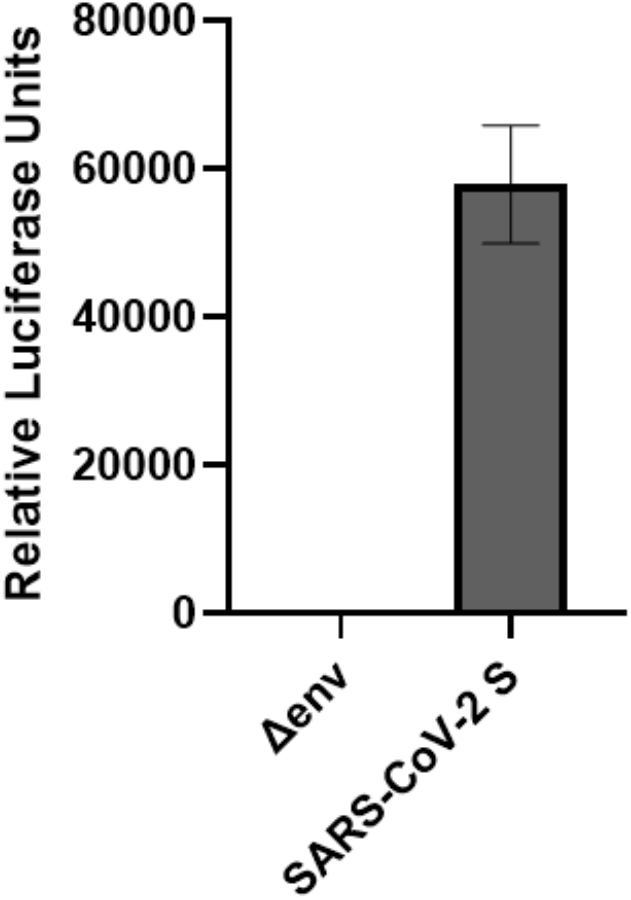
Characterization of the MLVpp system. Vero E6 cells were infected with MLV particles carrying the SARS-CoV-2 S protein or no envelope protein (negative control) and assessed for luciferase activity. The data shown is representative of the particle infectivity and error bars are standard deviation of three technical replicates.

**Figure S3:**
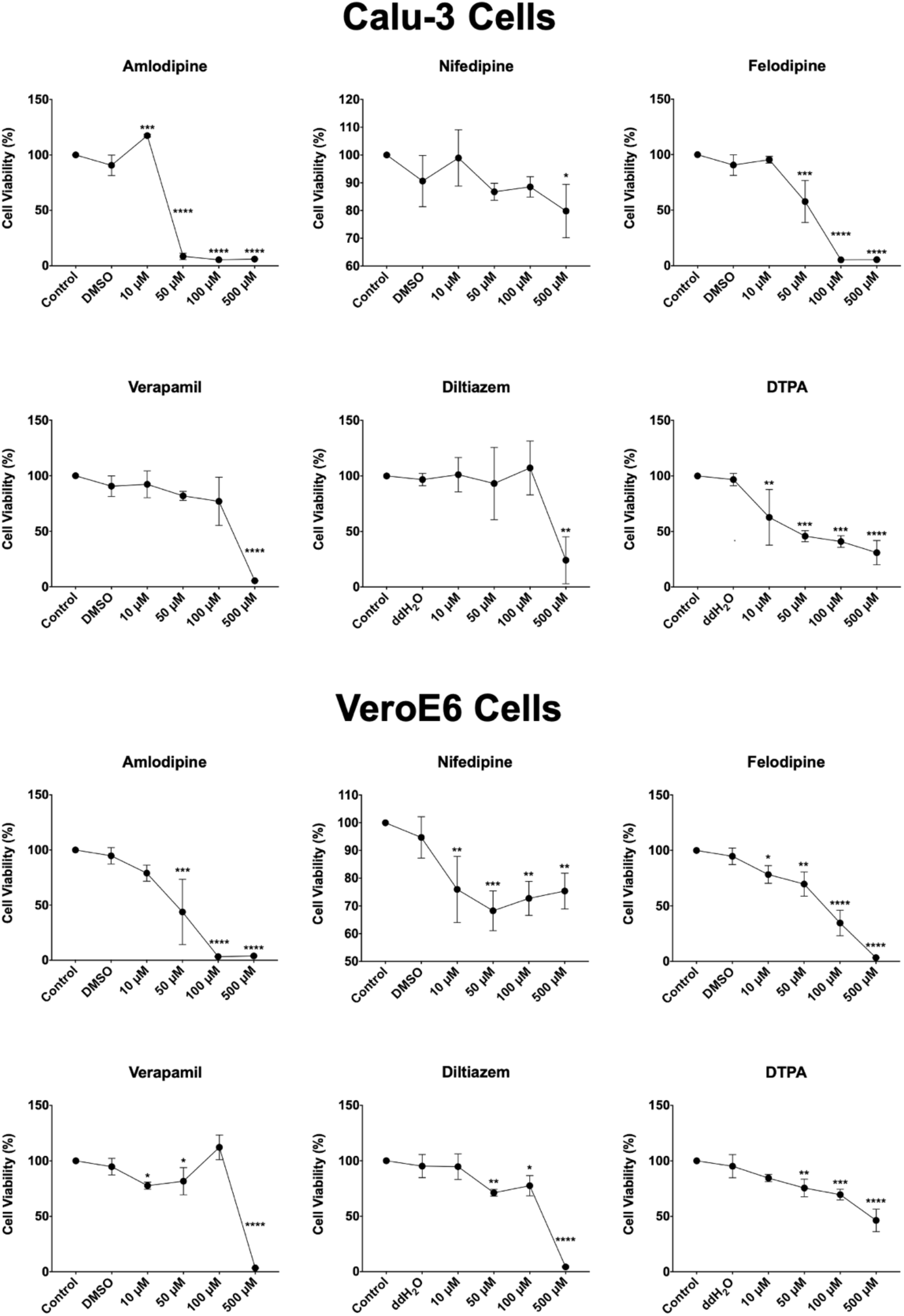
72-hour cell viability study. Vero E6 and Calu-3 cells were treated with the indicated concentrations of amlodipine, nifedipine, felodipine, verapamil, diltiazem and DTPA for 72 hours. After 72 hours, cell viability was measured using 3-(4,5-dimethylthiazol-2-yl)-2,5-diphenyltetrazolium bromide (MTT). Cell viability was determined by normalizing absorbance from the sample well by the average absorbance of untreated wells. As for the infections, 0 µM sample contained only DMSO or water depending on the solvent used for the drugs. Error bars represent standard deviations (n = 3). Asterisks indicate statistical significance compared to the untreated control. Statistical analysis was performed using an unpaired Student’s t-test. ^*^ = P < 0.05, ^**^ = P < 0.01, ^***^ = P < 0.001.

